# Integrating Chromosome Conformation and DNA Repair in a Computational Framework to Assess Cell Radiosensitivity

**DOI:** 10.1101/2024.03.25.586626

**Authors:** Matthew Andriotty, C.-K. Chris Wang, Anuj Kapadia, Rachel McCord, Greeshma Agasthya

## Abstract

**Objective:** The arrangement of chromosomes in the cell nucleus has implications for cell radiosensitivity. The development of new tools to utilize Hi-C chromosome conformation data in nanoscale radiation track structure simulations allows for *in silico* investigation of this phenomenon. We have developed a framework employing Hi-C-based cell nucleus models in Monte Carlo radiation simulations, in conjunction with mechanistic models of DNA repair, to predict not only the initial radiation-induced DNA damage, but also the repair outcomes resulting from this damage, allowing us to investigate the role chromosome conformation plays in the biological outcome of radiation exposure.

**Approach:** In this study, we used this framework to generate cell nucleus models based on Hi-C data from fibroblast and lymphoblastoid cells and explore the effects of cell type-specific chromosome structure on radiation response. The models were used to simulate external beam irradiation including DNA damage and subsequent DNA repair. The kinetics of the simulated DNA repair were compared with previous results.

**Main Results:** We found that the fibroblast models resulted in a higher rate of inter-chromosome misrepair than the lymphoblastoid model, despite having similar amounts of initial DNA damage and total misrepairs for each irradiation scenario.

**Significance:** This framework represents a step forward in radiobiological modeling and simulation allowing for more realistic investigation of radiosensitivity in different types of cells.

## INTRODUCTION

Recent advances in radiobiological modeling and simulation have produced several tools to predict the biological effects of radiation at the cellular scale. This includes accessible, open-source toolkits for nano-scale radiation track-structure simulation^1,2,3,4^ and mechanistic models of DNA repair.^2,5,6^ These tools are used to elucidate the various factors that affect cell radiosensitivity and cellular response to radiation in a variety of scenarios, taking into account the effects of radiation type, energy, and dose rate. However, they do not account for all factors that affect cell response to radiation.

One such factor is the three-dimensional arrangement of chromosomes in the cell nucleus, which has been shown in experimental studies to influence DNA damage and repair outcomes following radiation exposure.^7,8,9,10^ For example, misrepair of double-strand breaks (DSBs)–i.e., two broken ends of DNA that the cell attempts to reconnect–results in chromosome aberrations that could make cell division impossible, subsequently resulting in cell death or an increase in the likelihood of carcinogenesis. Therefore, the induction and repair of DSBs is of great importance in cellular radiobiology research.

If a cell’s DNA shows a high degree of clustering between certain parts of the genome, then DSBs occurring in that region will show a similar level of clustering. The proximity of the DSBs broken ends has implications on how they will be repaired. Regions with higher concentrations of broken ends will show increased probability of being incorrectly reconnected by the cell’s repair mechanisms,^8^ subsequently increasing the likelihood of chromosome aberrations. Therefore, the effect that clustering of genome structure or densely packed DNA (which may require chromatin decondensation and motion of DSBs to facilitate repair^11^) has on DNA damage and repair consequent to radiation exposure must be considered in simulations of cellular radiobiology.

Various studies have simulated radiation effects in models of whole cell nuclei. TOPAS-nBio^4^ and MEDRAS-MC^6^ have been used to simulate DNA damage and repair following proton irradiation in a model of a spherical human cell nucleus uniformly filled with DNA;^12^ however, that model does not account for variations in nucleus shape or arrangement of chromosomes. Others have also coupled radiation track structure simulations with DNA models to simulate DNA damage and repair outcomes, including models that are cell-type specific with regions of different chromatin compaction.^13,14,15,16^ The DNA structures in those models are constructed mathematically; they do not represent specific chromosome structures observed in certain populations of cells. In order to better represent chromosome structures of specific types of cells, Ingram, et al.^17^ developed a method using three-dimensional models based on Hi-C data in Geant4-DNA radiation track-structure simulations to predict initial DNA damage– but not repair outcomes–in these models.

To further investigate the effect of chromosome conformation on radiation-induced damage and repair, we leveraged tools for creating three-dimensional models of cell nuclei based on chromosome conformation data from real-world cells for use in TOPAS-nBio^4^ radiation track-structure simulations. The data used to create unique cell nucleus models comes from the Hi-C method, a type of chromosome conformation capture technology.^18^ This method results in an interaction matrix that represents the frequency of contacts within the 3D chromosome structure. Unlike other chromosome conformation capture methods, Hi-C data are ideal for creating three-dimensional models of whole cell nuclei, because they test interactions between all possible pairs of genome fragments, allowing us to infer a genome-wide arrangement of chromosomes. These data can be analyzed to determine topologically associating domains (TADs),^19^ which are regions of the genome in which segments of DNA show significantly greater contact with each other than with DNA outside the TAD. Once the genome is segmented into TADs, this information is used to create a three-dimensional representation of the genome made up of units representing TADs and organized according to contact probabilities between TADs.^17^

Three-dimensional chromosome structures follow patterns that are cell-type specific. This phenomenon gives Hi-C data utility for *in silico* studies of radiosensitivity between cell types. The unique TAD features and their arrangement in the nucleus of a specific type of cell can be used to create a model that represents chromosome contact frequencies in that cell type, and therefore can be compared with other types of cells in a radiation track-structure simulation.

The purpose of this work was to develop a computational framework that enables the study of the influence of DNA 3D structure and proximity of DSBs on biological outcomes of the DNA DSBs and repair processes. In a step forward for cellular-level radiobiological simulations, we combined radiation track-structure simulations in Hi-C-based cell nucleus models with mechanistic simulations of DNA repair. This allowed us to investigate the effect of varying chromosome structure on not only the initial DNA damage following irradiation, but also the yield of different types of DNA misrepairs, which is directly related to the biological outcome for the cell. We used Hi-C data from a study of X-ray irradiation in different types of cells^10^ as inputs for our simulation framework.

## METHODS AND MATERIALS

The simulation framework (Figure 1) developed in this work includes three main parts: (1) a method for generating three-dimensional models of cell nuclei from raw Hi-C data, (2) simulation of radiation track structure and DNA damage in these models, and (3) simulation of DNA repair to predict biological outcomes. The Hi-C data used in this work came from an *in vitro* study of BJ-5ta fibroblasts and GM12878 lymphoblastoid cells and their response to X-ray irradiation.^10^ Results from the *in vitro* study were compared to our simulations as validation for the developed framework.

**Figure 1.**
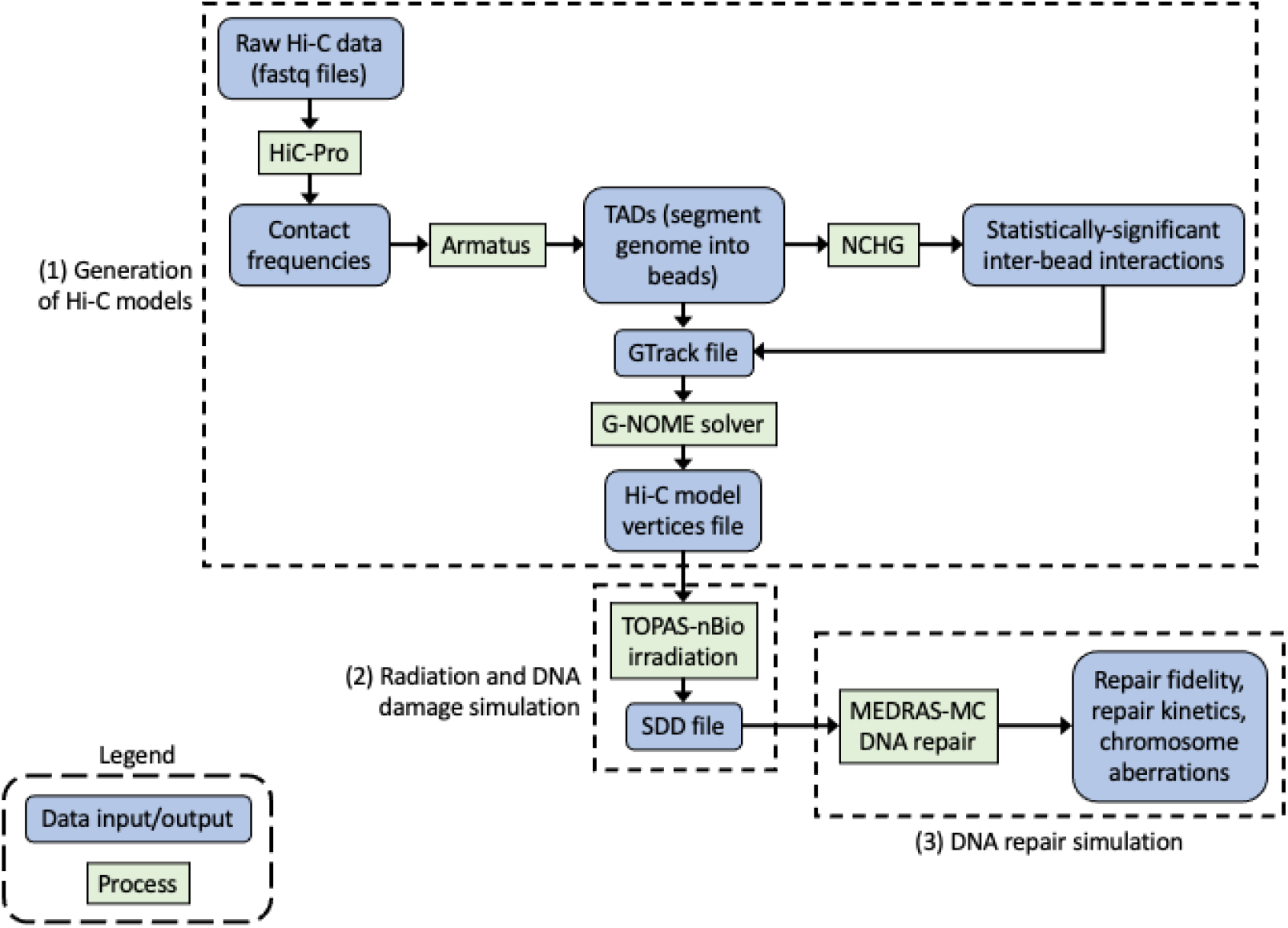
Flowchart of simulation framework developed to study the effects of chromosome structure on the outcomes of radiation-induced DNA damage and repair. The blue boxes represent data input/output steps and the green boxes represent different processes simulated as part of the framework. The dotted line boxes represent the three main parts of the framework: (1) generation of cell nucleus models from Hi-C data, (2) simulation of radiation and initial DNA damage, and (3) simulation of the repair of this DNA damage.

### Generation of cell nucleus models from Hi-C data

The method for processing raw Hi-C data to create cell nucleus models was adapted from work done by Ingram, et al.^17^ HiC-Pro^20^ was used to process the raw Hi-C data (in fastq format) to determine the raw inter- and intra-chromosomal contact frequencies. This information then served as input for Armatus,^21^ an algorithm for calling TADs. The number and locations of the TADs are a property of the type of cell as calculated from the Hi-C data. The TAD data were used to partition the entire genome into beads, each of which represents a TAD. The statistically significant inter-bead interactions were determined by the non-central hypergeometric distribution (NCHG). This information was arranged into a “gtrack” file containing the position of each bead within the genome, their sizes, and the inter-bead interactions. The size of the bead represents the number of base pairs contained within each TAD-defined segment. The G-NOME (Nuclear Organisation Modelling Environment) solver^17^ described by Ingram, et al. used the gtrack file to produce models in which spherical beads representing TADs were arranged according to their inter- and intra-chromosomal contacts. The output of the G-NOME solver was a file containing the spatial coordinates of each bead, the chromosome to which it belonged, and its radius.

### Radiation track-structure and DNA damage simulation

The TOPAS-nBio toolkit^4^ was used to simulate the irradiation of the cell nucleus models. The output files generated by the G-NOME solver were used to create spherical geometry components in TOPAS-nBio to represent each bead.

TOPAS-nBio is capable of running Monte Carlo simulations of the interactions of radiation with water at the nanoscale. The energy depositions from primary and secondary particles were recorded. If energy depositions within a range of 5 to 37.5 eV were recorded within one of the TADs (beads) in the cell nucleus model, then there was a probability–corresponding to the sensitive fraction of the bead–of a single-strand break (SSB) being scored at that location. The sensitive fraction represents the fraction of the volume of each bead in which energy depositions may cause damage to the sugar-phosphate backbone. The probability increases from zero at 5 eV to its maximum at 37.5 eV. SSBs were randomly assigned to one of the DNA strands with equal probability, and SSBs on opposite DNA strands within 3.2 nm of each other were scored as DSBs. Each bead was assigned to a particular chromosome, and the size of the bead represented the number of base pairs contained within. Using this information, a strand break’s position along the chromosome–in terms of genetic length–was determined by calculating the total number of base pairs in the chain of beads from the start of the short arm of the chromosome to the bead in which the strand break occurred.^17^ The locations of the DSBs (both spatial coordinates and location on the chromosome) were recorded in a Standard DNA Damage (SDD) file.^22^

The use of TOPAS-nBio in this framework allows the user to conveniently adjust the type, energy, and dose of radiation used in the simulation. Therefore, future studies could employ this framework in investigations of a wide variety of radiotherapy and radiation protection applications. In summary, at this step in the framework we can estimate the number and location of SSBs and DSBs as a consequence of interaction of radiation with a cell nucleus.

### Mechanistic DNA repair simulation

To predict the biological outcome of the above estimated radiation-induced DNA damage, MEDRAS-MC (Mechanistic DNA Repair and Survival-Monte Carlo) was used to simulate the repair of the DSBs.^6^ MEDRAS does not take SSBs into account, and they are assumed not to have consequences for the misrepair outcomes for the purpose of this work. The SDD files from TOPAS-nBio served as input for the MEDRAS code, which simulated the rejoining of the broken DSB ends by one of three different repair pathways: nonhomologous end joining (NHEJ), homologous recombination (HR), and microhomology mediated end joining (MMEJ). HR uses the sister chromatid (available after DNA replication) as a template to facilitate the repair of DSBs with low probability of error. In the event that HR is unavailable, NHEJ may be used to rejoin broken ends without a template, resulting in a greater error rate. If NHEJ is inhibited–as in cancer cells, for instance–MMEJ may be used as a highly error-prone alternative.^23^ MEDRAS chooses the repair pathway according to the complexity of the DSB as determined from the distribution of damage in the SDD file; simple breaks are repaired by NHEJ, while complex breaks are repaired by HR if sister chromatids are available and NHEJ if they are not, as is observed in repair-competent cells. MMEJ may be used if a DSB fails to be repaired by HR and NHEJ due to defects in those pathways, which is observed in repair-defective cells. MEDRAS uses a Monte Carlo method to rejoin DSBs by calculating the interaction rates between all pairs of broken ends as a function of distance and the selected pathway’s likelihood of success.^6^

At the end of this step, the kinetics of DNA repair were calculated to obtain the number of unrepaired DSBs remaining as a function of time after irradiation. We also calculated the yield of DSB misrepairs and inter-chromosome aberrations in each cell model 24 hours after irradiation.

### Validation and experimentation with developed framework

Three cell models were developed using the framework described above: (1) spherical lymphoblastoid (radius = 5 µm), (2) spherical fibroblast (radius = 5 µm), and (3) ellipsoidal fibroblast (semi-axes = 11.181 µm, 11.181 µm, and 1 µm). The thickness of the flattened ellipsoid was chosen according to the thickness of human fibroblast cells reported in experimental studies.^24^ The sizes of the other semi-axes of the ellipsoid were chosen to constrain the beads within an envelope of equal volume to that of the spherical nuclei. The G-NOME solver was run for two million iterations to obtain optimized solutions for the geometry of spherical nucleus models using data from both fibroblast and lymphoblastoid cells.^10^ Additionally, the solver was run for four million iterations with the fibroblast data to create a flattened ellipsoid nucleus in order to more accurately reproduce the *in vitro* geometry of that type of cell. A greater number of iterations was used for the ellipsoidal model to ensure that beads were properly constrained within the ellipsoidal envelope.^17^

For this work, a 5-Gray irradiation with 160 kVp X-rays was simulated for each cell model in TOPAS-nBio for comparison with the *in vitro* experiment from which the Hi-C data were sourced. SpekPy^25^ was used to estimate the X-ray spectrum, and a TOPAS simulation was used to estimate the energies of secondary electrons resulting from the X-ray spectrum. The secondary electron spectrum–rather than the X-rays themselves–was simulated directly in TOPAS-nBio for more efficient computation. The numbers of DSBs after irradiation with each radiation source were estimated and the corresponding SDD files were used to calculate outcomes from the repair processes.

The repair kinetics following X-ray irradiation in the three cell models were simulated using MEDRAS, and the outcomes were compared with *in vitro* data (the γH2AX intensity 30 minutes and 24 hours after irradiation). Additionally, the yields of DSB misrepairs and inter-chromosome aberrations in the three cell models were calculated.

Additional experiments were conducted to evaluate the versatility of this framework and compare the effects of different types and energies of radiation in these cell models. Both 1-MeV and 100-MeV protons were simulated and the yield of DSB misrepairs in all three cell models were calculated. MEDRAS was also used to calculate the yield of different types of chromosome aberrations in the cell nucleus models and visualize these results in mFISH plots to illustrate the different possible outcomes of DNA misrepair. The outcomes of the validation and experiments are shown in the results section. Welch’s *t*-test was used to compare simulated and *in vitro* DNA repair kinetics. One-way ANOVA was used to compare repair outcomes between the cell models for each type of radiation simulated.

## RESULTS

### Analysis of the cell nucleus models

Cell nucleus models were simulated for the GM12878 lymphoblastoid and BJ-5ta fibroblast cells. A spherical model was simulated for each cell type (Figures 2 and 3a), and an additional flattened nucleus model (Figure 3b) was simulated for the fibroblasts. The number of TADs (beads) in the fibroblast models was 8,696 for both the spherical and flattened cells. In contrast, the lymphoblastoid models contained 10,102 TADs (beads). The number of beads is determined by the TADs found when processing the Hi-C data.

**Figure 2.**
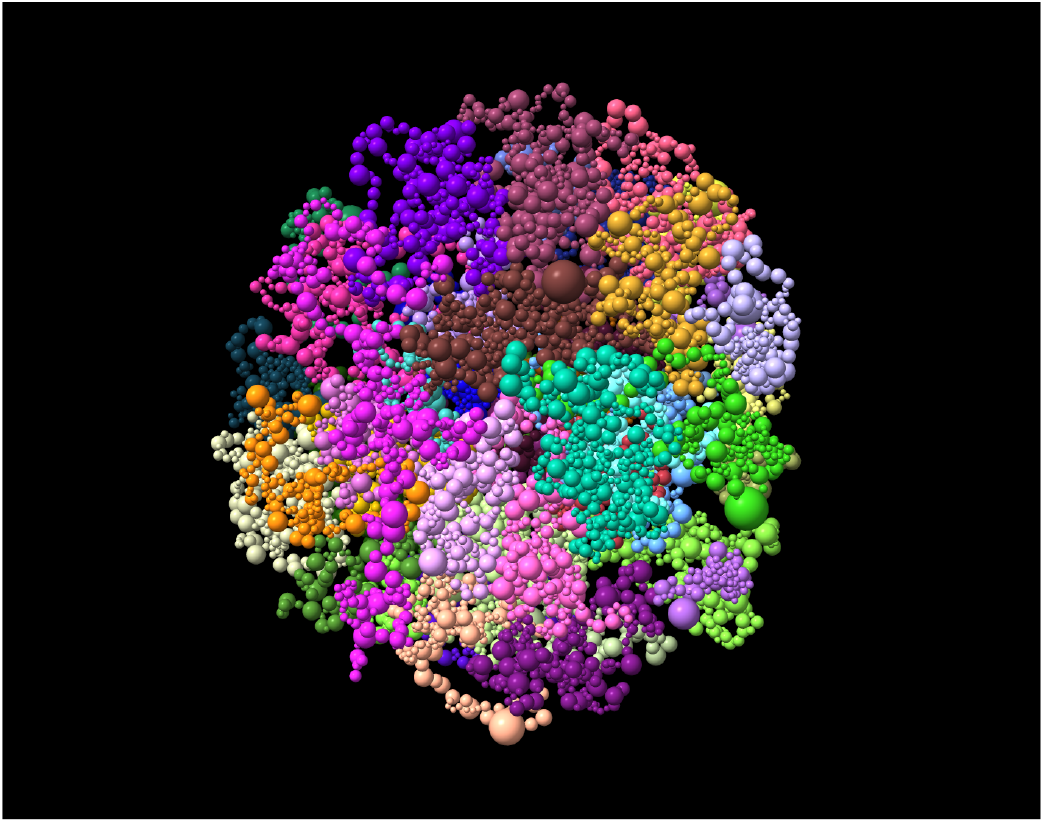
GM12878 lymphoblastoid model created using G-NOME and visualized using ChimeraX. Each bead represents a TAD, and the size of the bead represents the number of base pairs contained within. The color of each bead represents the chromosome to which it belongs.

**Figure 3.**
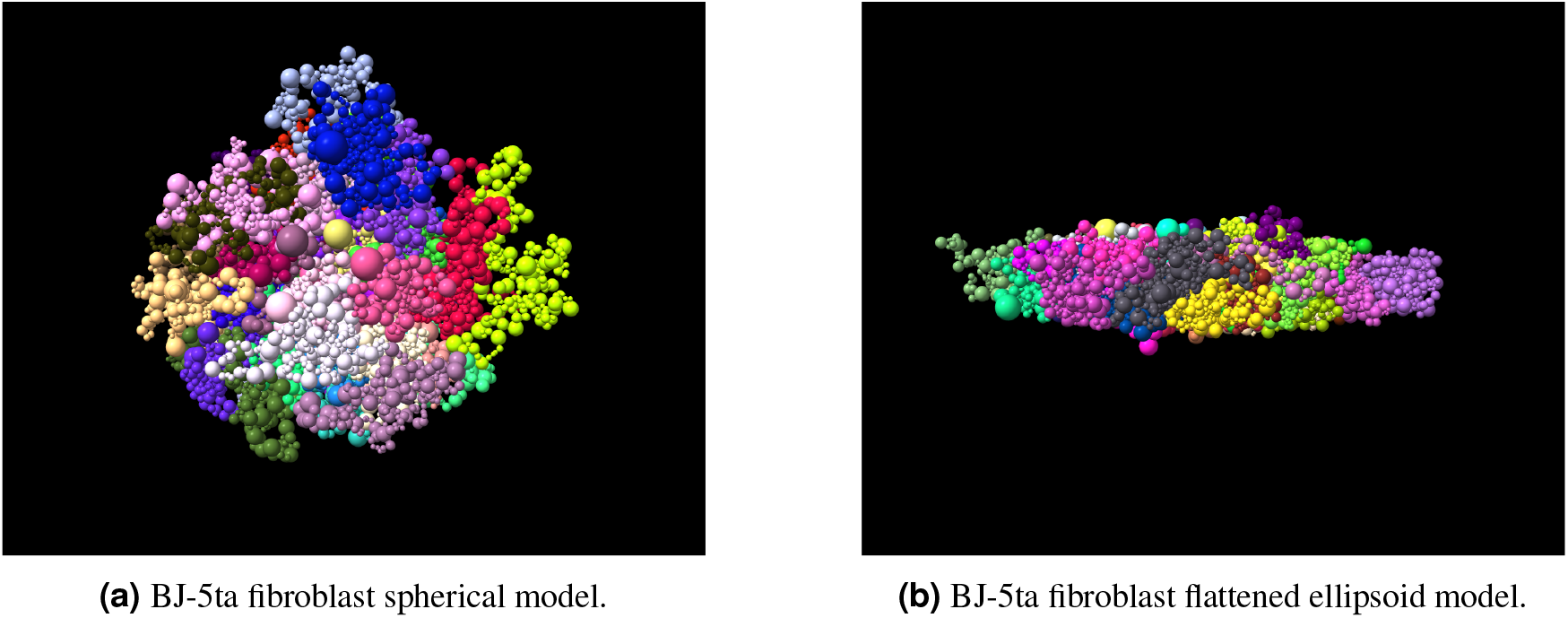
BJ-5ta fibroblast models: (a) spherical, and (b) flattened ellipsoid. Each bead represents a TAD, and the size of the bead represents the number of base pairs contained within. The color of each bead represents the chromosome it belongs to.

The radial distribution of TADs (beads) was calculated in order to evaluate the differences between cell models. A histogram of the distances of TADs from the center of the nucleus for all three cell nucleus models (Figure 4) shows that the lymphoblastoid model has a higher concentration of TADs near the periphery of the nucleus than the spherical fibroblast model. The flattened fibroblast model had a greater width than the other two models (Figure 4a), but if the radial distances of the TADs are normalized for each model, the flattened fibroblast model has TADs that are more centrally located than the other two models’ (Figure 4b).

**Figure 4.**
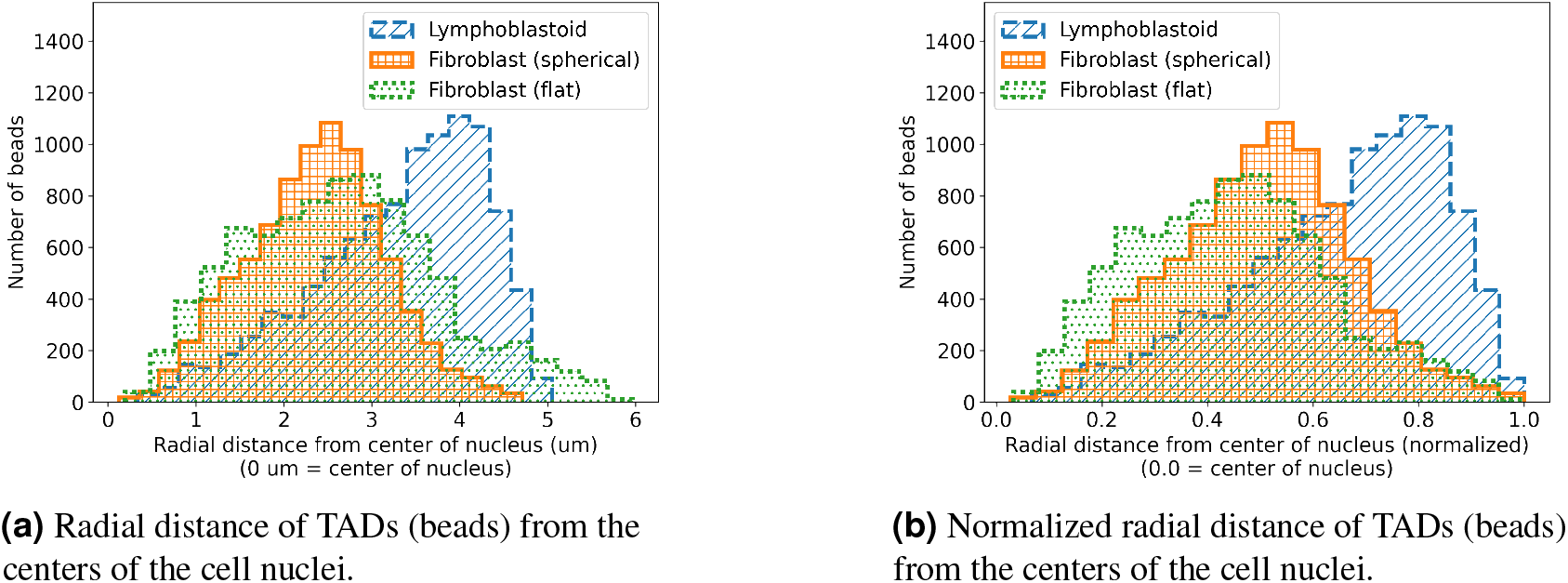
Comparison of the radial distances of TADs (beads) from the center of the nucleus across three cell nucleus models.

### Initial DNA damage

TOPAS-nBio was used to simulate the irradiation of these cell models and record the resulting DSBs. The yield of initial DSBs per Gray following irradiation was calculated for each cell model (Figure 5 and Table 1).

**Figure 5.**
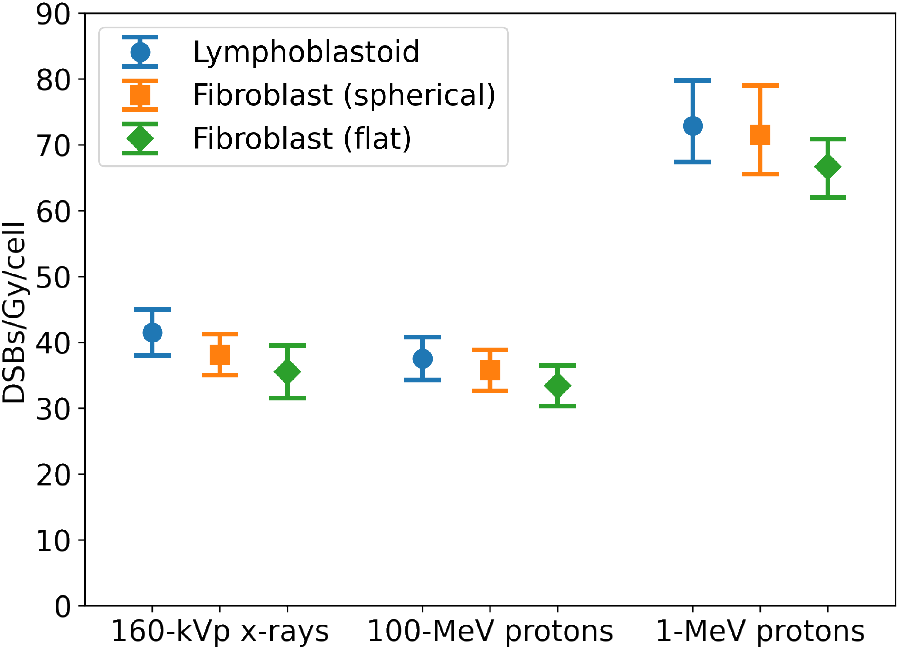
Yield of initial DSBs/Gy/cell post-irradiation in the three cell models (mean *±* SE).

**Table 1.** Yield of initial DSBs/Gy/cell post-irradiation in the three cell models (mean *±* SE).

|  | Lymphoblastoid | Fibroblast (spherical) | Fibroblast (flat) |
| --- | --- | --- | --- |
| 160-kVp X-rays | 41.5 ± 3.5 | 38.1 ± 3.1 | 35.6 ± 4.0 |
| 100-MeV protons | 37.5 ± 3.2 | 35.7 ± 3.5 | 33.4 ± 3.1 |
| 1-MeV protons | 73.0 ± 5.8 | 71.3 ± 6.1 | 66.7 ± 4.6 |

### DNA repair kinetics

After simulating the irradiation of these cells in TOPAS-nBio, MEDRAS was used to simulate the kinetics of DNA repair. For each cell model, the vast majority of DSBs were repaired in the first few hours after irradiation, which is consistent with previous literature.^26^

Figure 6 shows the ratio of DSB repair foci remaining 30 minutes to 24 hours after irradiation. This was compared to *in vitro* data from the study that generated this Hi-C data.^10^ In that work, the γH2AX intensity 30 minutes and 24 hours after 160 kVp irradiation was reported. A *t*-test for comparison of means between simulated and experimental data showed that the simulated repair foci ratio is not significantly different from the the experimental ratio in the two fibroblast models, but the lymphoblast model showed a significant difference (Table 2). All simulated means are within one standard deviation of the experimental means.

**Figure 6.**
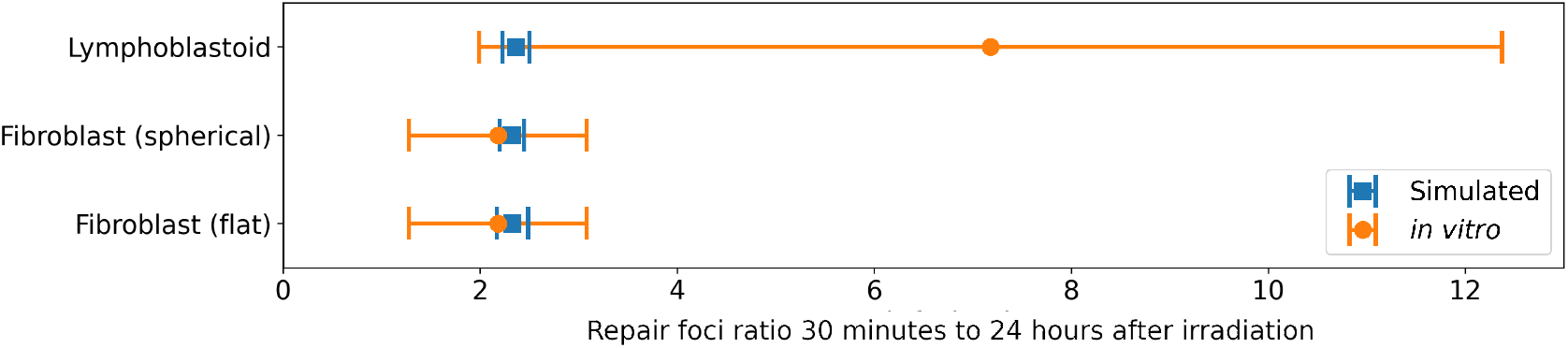
Ratio of γH2AX foci relative to baseline 30 minutes to 24 hours after 160-kVp X-ray irradiation and comparision with *in vitro* data (mean *±* SD). Comparison of means between simulated and *in vitro* data: Lymphoblastoid: *p <* 0.001; Fibroblast (spherical): *p* = 0.34; Fibroblast (flat): *p* = 0.31.

**Table 2.** Results of a Welch’s *t*-test comparing the means between *in silico* and *in vitro* data for each cell type (significance level: 0.05).

|  | Lymphoblastoid | Fibroblast (spherical) | Fibroblast (flat) |
| --- | --- | --- | --- |
| *p*-value | < 0.001 | 0.34 | 0.31 |

### DNA repair fidelity

A plot of misrepair yield per Gray (Figure 7) shows the likelihood of a DSB being misrepaired in each cell model after 160-kVp X-ray irradiation. The lymphoblastoid model showed a slightly greater probability of misrepair than the spherical fibroblast model, which yielded slightly more misrepairs than the flattened fibroblast model, the same pattern observed with initial DSBs.

**Figure 7.**
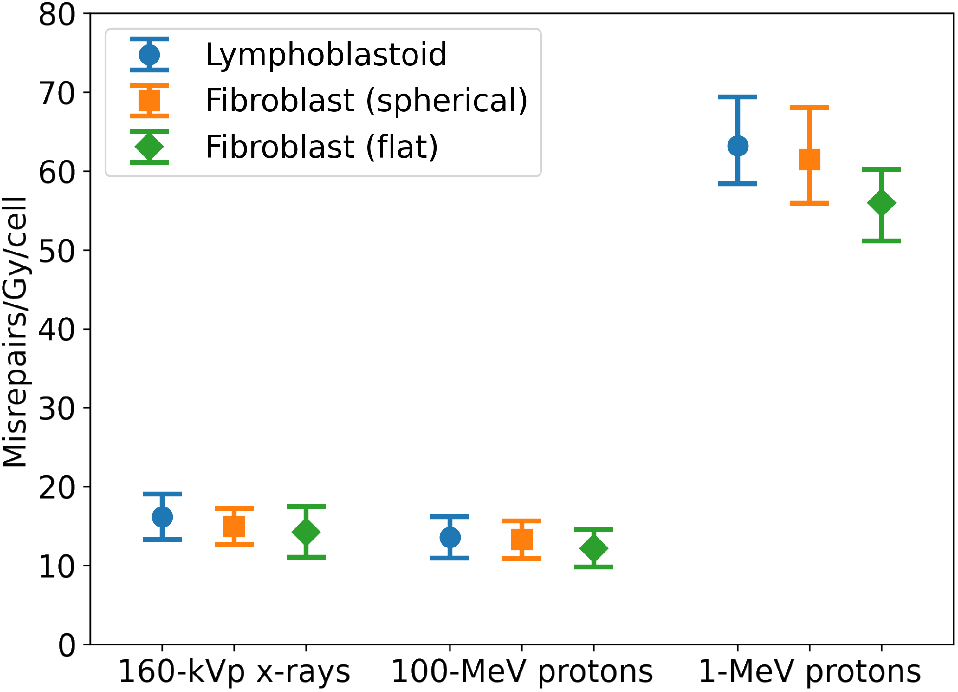
Misrepair yields following irradiation and repair (mean *±* SE).

For comparison between different types of radiation, 100-MeV and 1-MeV protons were simulated in TOPAS-nBio as well. The relative misrepair probabilities between cell types were similar to the X-ray results (Figure 7). The 1-MeV proton irradiation, having a greater LET than the other types of radiation used in this work, resulted in a much greater yield of DSB misrepairs. The results of ANOVA for misrepairs from each type of radiation are shown in Table 3.

**Table 3.** The results of one-way ANOVA comparing the mean total misrepairs/Gy/cell for each type of radiation. The results of Tukey HSD post-hoc tests are shown only for comparisons that had a significant difference (significance level: 0.05).

|  | *p*-value | Post-hoc tests showing significant difference |
| --- | --- | --- |
| 160-kVp X-rays | 0.0033 | Lymphoblastoid vs flattened fibroblast: $p = 0.0027$ |
| 100-MeV protons | 0.0154 | Lymphoblastoid vs flattened fibroblast: $p = 0.0166$ |
| 1-MeV protons | 0.0153 | Lymphoblastoid vs flattened fibroblast: $p = 0.0156$ |

The yield of misrepairs involving DSBs from two different chromosomes is shown in figure 8. Contrary to the pattern shown for total misrepairs, the fibroblast model yielded more inter-chromosome misrepairs than the lymphoblastoid model, with the flattened fibroblast model yielding the greatest number of inter-chromosome misrepairs. Table 4 shows the results of ANOVA for inter-chromosome misrepairs for each type of radiation.

**Figure 8.**
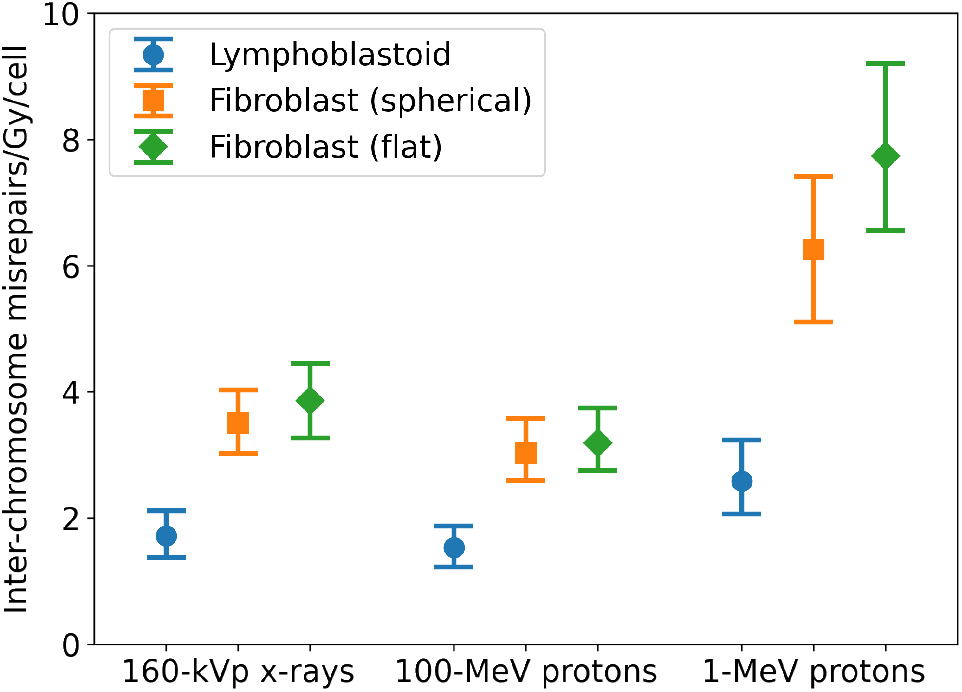
Inter-chromosome misrepair yields following irradiation and repair (mean *±* SE).

**Table 4.** The results of one-way ANOVA comparing the mean inter-chromosome misrepairs/Gy/cell for each type of radiation. The results of Tukey HSD post-hoc tests are shown only for comparisons that had a significant difference (significance level: 0.05).

|  | *p*-value | Post-hoc tests showing significant difference |
| --- | --- | --- |
| 160-kVp X-rays | < 0.001 | Lymphoblastoid vs either fibroblast: $p < 0.001$ |
| 100-MeV protons | < 0.001 | Lymphoblastoid vs either fibroblast: $p < 0.001$ |
| 1-MeV protons | < 0.001 | Lymphoblastoid vs either fibroblast: $p < 0.001$ |

To demonstrate prediction of specific types of chromosome aberrations as a consequence of DNA repair, two mFISH plots have been included (Figure 9) illustrating the status of all 23 pairs of chromosomes in the fibroblast cell model following X-ray irradiation and DNA repair. Figure 9a shows an inter-chromosome aberration which involves the the rejoining of fragments from two different chromosomes. Figure 9b shows a ring aberration in which two ends of a fragment have been rejoined to each other rather than to the remaining part of the chromosome. Future work will leverage this capability to predict cell survival following radiation exposure by categorizing these types of aberrations into lethal and non-lethal.

**Figure 9.**
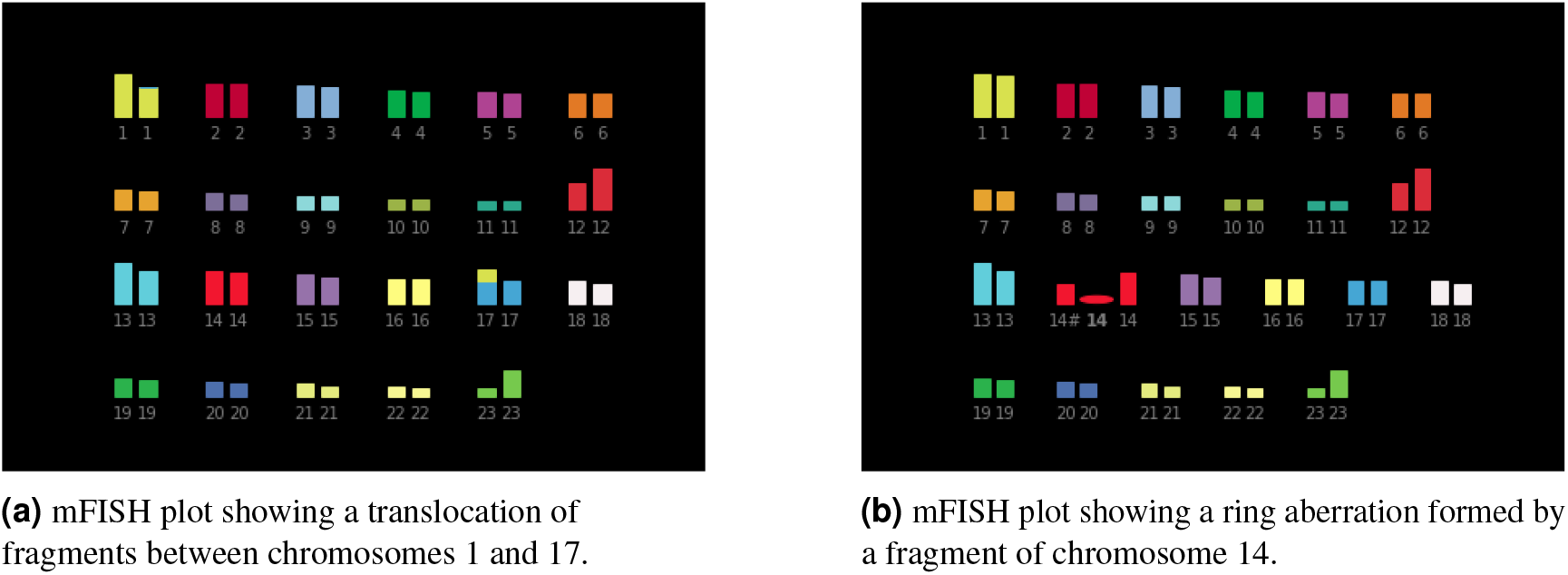
mFISH plots generated by MEDRAS simulations of DNA repair in fibroblast cell models following X-ray irradiation.

## DISCUSSION

The simulation framework developed in this study models varying and specific chromosome conformations from different cell nuclei and biological endpoints such as DSBs, SSBs, DSB misrepair and chromosome aberrations. The framework can model differences in radiation response corresponding to differences in chromosome structures of cells.

The differences in the geometric structure of the cell nucleus models developed for this work demonstrate that this method accurately captures and represents the cell-type-specific chromosome conformations from *in vitro* cells. These unique cell models resulted in different radiation damage repair outcomes between cell types, indicating that the chromosome structure variation influences not only the distribution of initial DSBs, but also the likelihood of certain types of misrepair, which is closely connected with the biological outcome (cell death or non-lethal mutation) of radiation exposure. The inclusion of DNA repair models in this framework and the demonstration that chromosome structure affects simulated DNA repair outcomes is an encouraging step forward in the use of these tools to investigate the factors that contribute to differences in cell radiosensitivity.

The validation of *in silico* methods should be considered in order to ensure that these methods are accurately producing results observed in real cells and to lend credibility to results obtained when applying these methods to new scenarios. The *in vitro* study that provided the Hi-C data for this work allowed for some comparison between simulated and experimental results. The kinetics of DNA DSB repair in the fibroblast models were shown to be consistent with the change in γH2AX intensity in the 24 hours following irradiation. One limitation of these simulations was that non-irradiated real world cells have a non-zero number of γH2AX foci at any given time. At present, our model does not account for foci that were not induced by radiation damage. Additionally, the repair simulations in this work did not take into account differences in repair capabilities between cell types; the only difference between cell types considered in this work was the difference in chromosome conformation based on Hi-C data. It is for this reason that the three cell models had similar *in silico* repair kinetics. Future work will include adjusting parameters of the repair simulation to include differences in the availability of DNA repair pathways.

More comprehensive, direct validations are desirable to further demonstrate the effectiveness of this framework for the prediction of post-irradiation biological outcomes. Future work includes coordination with *in vitro* experiments to more directly compare simulated and real-world metrics, such as the yield of specific types of chromosome aberrations or cell survival. These results would increase confidence in the validity of these models.

The observation of greater yields of inter-chromosome misrepairs in the fibroblast models–despite similar yields of initial DSBs and total misrepairs–is a remarkable result of the differences in chromosome conformation between these cell nucleus models. The radial distributions of TADs (beads) showed that the TADs tended to be more centrally located in the fibroblast models than in the lymphoblastoid models, which had more TADs located near the periphery of the nucleus. The increased proximity of TADs in the central region of the nucleus may be contributing to more inter-chromosome intermingling of TADs–and therefore DSBs–in the fibroblast models, leading to an increased likelihood of misrejoining of broken ends from different chromosomes.

## CONCLUSION

Because cell type, cell-cycle phase, and disease state are associated with differences in chromosome conformation, further development of this framework will provide insights into factors that contribute to variations in the biological response to radiotherapy. Capturing and modeling these differences will help model the efficacy of treatment at the cellular scale. Ongoing work includes the development of a multiscale *in silico* framework for precision radiation therapy dosimetry capable of modeling cell toxicity, radiosensitivity, and repair mechanisms, along with tumor growth models and biodistribution of radiation at the tissue, organ, and whole-body scales to predict treatment outcomes. The integration and interaction of the framework described in this work with the multiscale model will allow for highly adaptable and specific predictions of the biological outcomes of radiation exposure at multiple scales.

## ACKNOWLEDGMENTS

This work was supported by the Office of Biological and Environmental Research (BER) and Laboratory Directed Research and Development Program of Oak Ridge National Laboratory, managed by UT-Battelle, LLC, for the U.S. Department of Energy. This research used resources of the Compute and Data Environment for Science (CADES) at the Oak Ridge National Laboratory. This manuscript has been authored by UT-Battelle, LLC under Contract No. DE-AC05-00OR22725 with the U.S. Department of Energy.

